# Engineered bacterial M1GS ribozyme efficiently cleaves the most abundant ribosomal RNA in a human cancer cell line

**DOI:** 10.1101/2025.11.07.687305

**Authors:** Neha Priyadarshini, Navinchandra Venkatarama Puppala, Harikrishnan Poiyamozhi, Swati Biswas, Gireesha Mohannath

## Abstract

M1GS, an engineered Ribonuclease P (RNase P), can be customized to specifically target and cleave any RNA. It involves tethering an *Escherichia coli* M1 RNA to a short guide sequence (GS) complementary to the RNA targeted for cleavage. We have developed a Python script-based bioinformatic tool built based on the requirements of M1GS, coupled to secondary structure predictions, to identify M1GS target sites for any given RNA. Primarily, we demonstrate the utility of the tool for predicting M1GS target sites across multiple RNAs, including human 28S ribosomal RNA (rRNA), and then we demonstrate efficient M1GS-mediated downregulation of 28S rRNA in a human cancer cell line. Efficient cleavage of rRNA, the most abundant RNA molecule, suggests a high turnover number for the M1GS ribozyme. Lastly, we discuss the utility of M1GS ribozyme-mediated rRNA downregulation as a potential anticancer modality in cancers where rRNAs are upregulated.

## Introduction

Targeted RNA cleavage has commonly been employed as a reverse-genetic tool to understand genes’ functions and as a defense modality against pathogens, including viruses. Commonly used tools include RNA interference (RNAi)/antisense RNA [1-4], engineered ribozymes [5], and recently developed CRISPR-Cas-based genome editing tools [6, 7].

Ribonuclease P (RNase P) RNA is among the earliest discovered ribozymes and is a ribonucleoprotein (RNP) complex found in all three domains of life [8, 9]. Although RNase P has a shared function in the 5’ tRNA maturation, the composition of the RNase P RNP complex is evolutionarily divergent [10-15]. In *Escherichia coli*, RNase P is made up of a catalytic RNA component called M1 RNA and a protein cofactor called C5 [16-18]. M1 RNA possesses catalytic activity *in vitro* in the absence of a protein cofactor [19]. RNase P recognizes precursor tRNA based on its structure, not the tRNA sequence. This feature was explored to engineer RNase P to selectively target and cleave any RNA molecule as a gene knockdown strategy [9, 20-22].

M1GS is an RNase P-based gene knockdown strategy, wherein *E. coli* M1 RNA is fused through its 3’ end to a guide sequence (GS), which is complementary to the target RNA. When the customized M1GS is expressed in the transfected cells, the GS hybridizes to the complementary sequence in the target RNA which is then cleaved by the tethered M1 RNA immediately upstream of the binding site (Figure 1) [15, 23, 24]. The M1GS strategy has been used to downregulate genes in both prokaryotes and eukaryotes, demonstrating its utility as an effective tool for disrupting genes’ functions [25-27]. Moreover, the M1GS strategy is simple, consisting of the expression of a single exogenous M1 RNA fused to a few short sequences. Its total length, including all the sequences expressed, is ∼590 nt. Further, it can be expressed using various types of inducible and constitutive promoters [28, 29]. An inactive M1GS carrying an enzymatically deficient mutant version of the M1 RNA has been included in multiple studies as a control to demonstrate that the mutant phenotype observed in their studies is due to the enzymatically active M1GS-mediated cleavage, not due to the GS-target RNA base-pairing or any other random effects of the M1GS expression [23, 26, 27, 30]. However, despite its simplicity, specificity, and effectiveness in cleaving targeted RNA, the M1GS strategy appears to be underutilized. Perhaps one reason is that prior knowledge of the M1GS target site requirements is necessary for its use, discouraging its frequent use.

**Figure 1:**
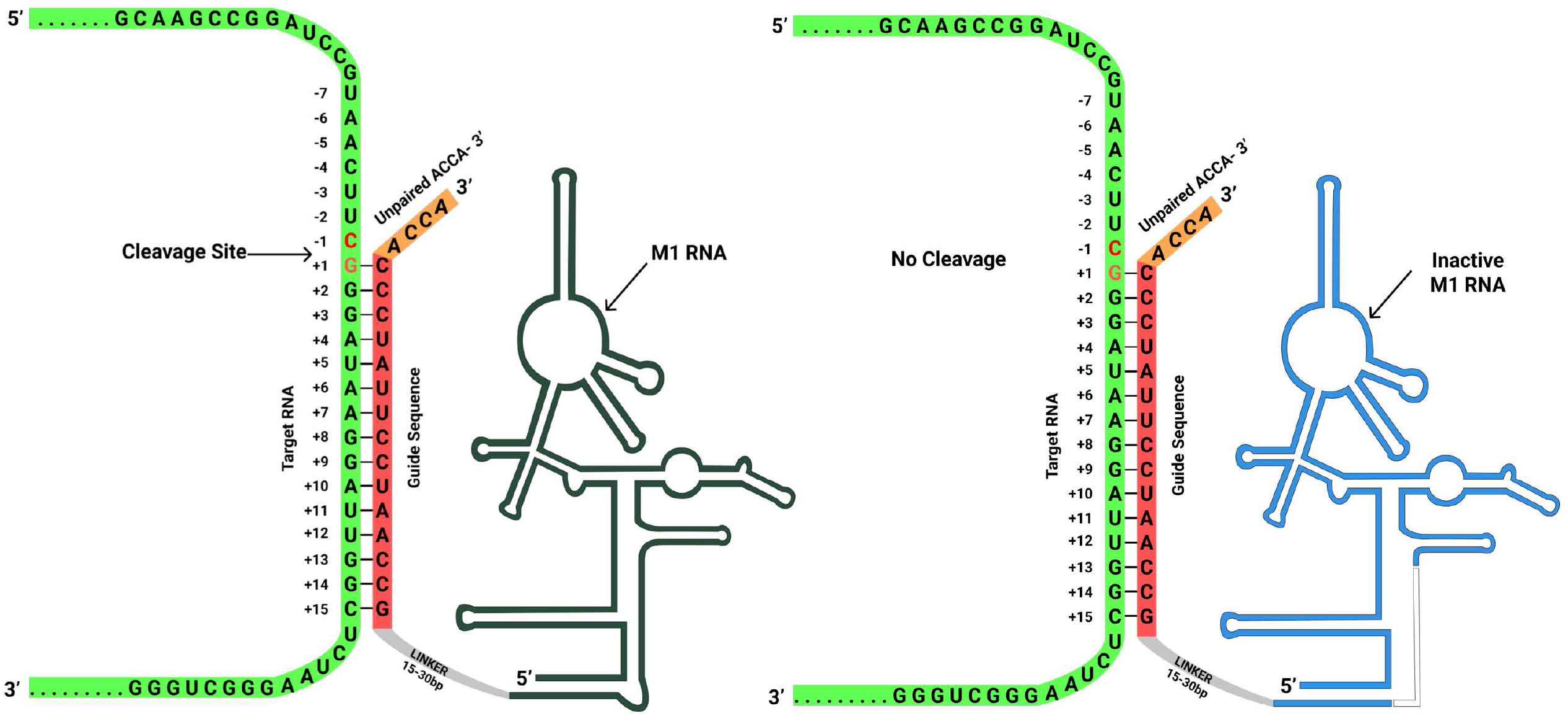
A schematic representation of the M1GS strategy to cleave a target RNA. The cartoon shows how the engineered RNase P ribozyme M1GS cleaves the target RNA by hybridizing to it through a complementary guide sequence (GS). Human 285 rRNA as a target RNA and the corresponding complementary guide sequence (GS) are shown in the diagram. The on the left represents M1GS with an active M1 RNA and the one on the right represents M1GS with an inactive, mutant M1 RNA.

Although M1GS has been employed to cleave multiple RNAs in human cells, it has not been tested to cleave ribosomal RNA (rRNA), the most abundant RNA species (up to 80% of total RNA). In the nucleolus, RNA polymerase I (RNA Pol I) transcribes 45S rRNA genes to produce pre-45S rRNA, which is then processed to 18S, 5.8S, and 28S rRNAs, which form the catalytic core of ribosomes, the protein-synthesising cell organelles (Figure S1). 45S ribosomal RNA genes are upregulated in several cancers [31-33], and therefore, RNA Pol I transcription has been therapeutically targeted in these cancers [34-40].

In this study, we describe *M1GS Target Site Finder*, a Python script-based user-friendly bioinformatic tool we developed to identify M1GS target sites for any given RNA using either DNA or RNA sequence as input. We demonstrate the utility of this bioinformatic tool in predicting M1GS target sites for multiple RNAs, including human 28S rRNA. Then, we demonstrate its downregulation via targeted cleavage by a customized M1GS ribozyme in a lung cancer cell line, using one of the GSs identified by the tool. Lastly, we discuss the potential advantages of M1GS ribozyme and M1GS ribozyme-mediated rRNA downregulation as a potential anticancer modality in cancers where rRNAs are upregulated.

## Results

We developed a Python script-based bioinformatic tool with a user-friendly web interface for automated identification of all possible M1GS cleavage sites for any target RNA. It provides the flexibility to customize the need-based predictions with various input options. A flowchart depicting the workflow of the *M1GS target finder* tool is shown in Figure 2. The tool can be accessed using the link (http://m1gstargetfinder.bits-hyderabad.ac.in:8000/m1gs/). Several input options that an end user can provide are described below. All the recommended input options are for M1GS involving *E. coli* M1 RNA only. Users have to determine if these options work equally well for other RNase P RNA.

**Figure 2.**
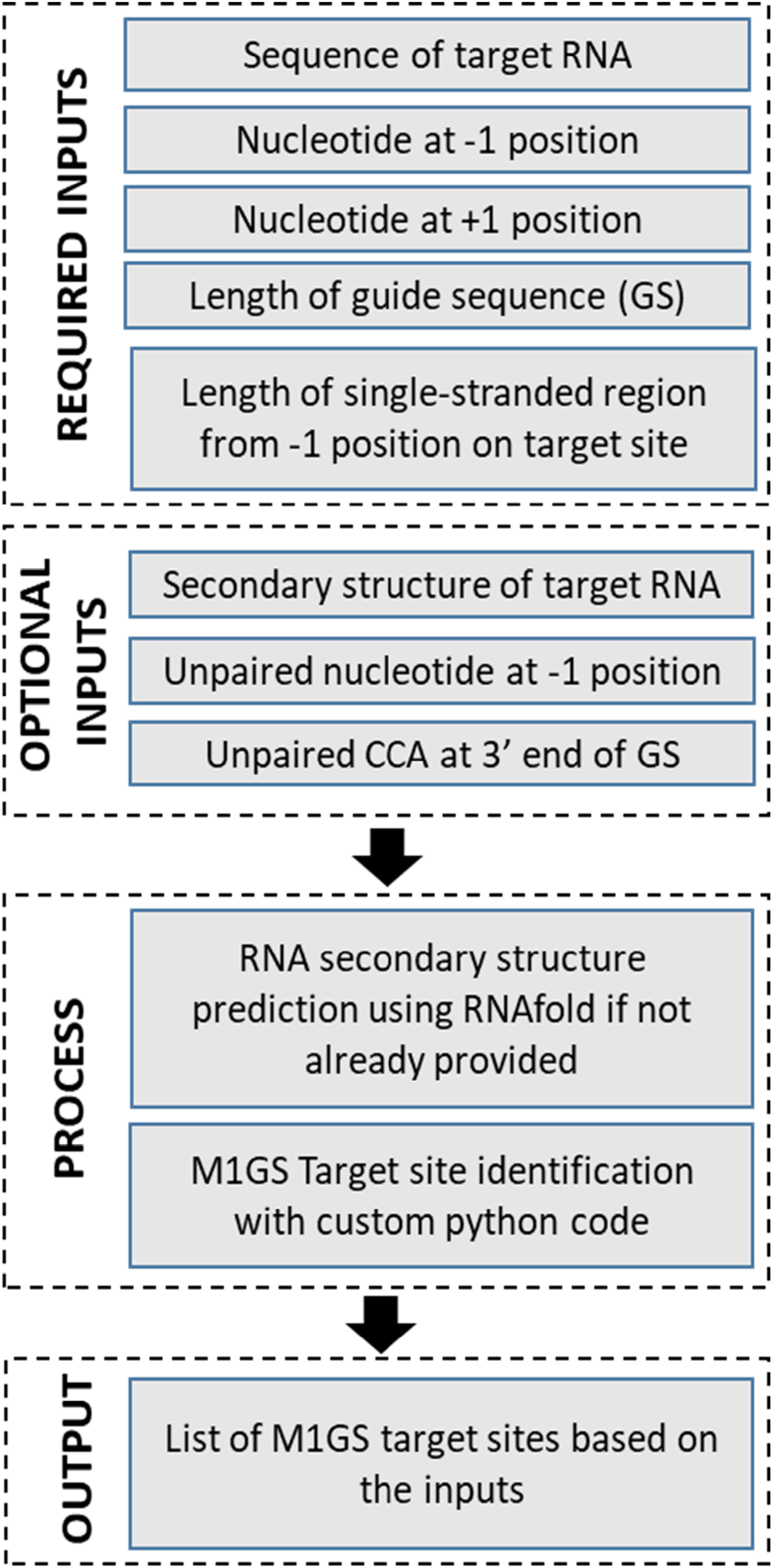
The flowchart depicts the operational modality of the *M1GS Target site finder* tool.

### Inputs for *M1GS target site finder* tool

#### Sequence of the target RNA

To identify all M1GS sites in a target RNA, as the first step, the sequence of the target RNA or DNA should be entered in the given space in the tool where it asks for ‘Sequences of target RNA/DNA’. If users provide a DNA sequence as input, the tool will automatically convert it to the corresponding RNA sequence and then use it to identify M1GS target sites.

#### The nucleotide at the -1 position

The 5’ nucleotide with respect to the M1GS cleavage site (-1 position) is a pyrimidine [15, 24]. Uracil is a conserved preference at the -1 position of the pre-tRNA in bacteria and archaea [41]. Consistent with this conservation, U at the -1 position was found to be critical for cleavage site selection and cleavage efficiency of tRNA substrates [42]. However, some bacterial species also have adenine, cytosine, or guanine at the -1 position, demonstrating that cleavage is also possible with nucleotides other than uracil at -1. Several studies have demonstrated customized M1GS-mediated cleavage of the target mRNA with varying nucleotides at the -1 position [43-45]. Therefore, we designed our tool to provide a choice of all four nucleotides at the -1 position. End users are recommended to prefer uracil at the -1 position, but if it is unavailable, they may choose cytosine (the other pyrimidine) or any other nucleotides of their choice.

#### The nucleotide at the +1 position

The identity of the base at the +1 position is a salient determinant for the M1GS cleavage site selection. A purine at the +1 position is important for both cleavage efficiency and cleavage site selection (between the -1 and +1 sites), regardless of whether the +1 nucleotide is base-paired or not [24]. When a purine at the +1 position is base-paired, it results in optimal cleavage of the target RNA [24]. In contrast, when a pyrimidine is at the +1 position and is not base-paired, it results in an aberrant cleavage [24]. Moreover, cleavage of a substrate with a purine at the +1 position is more efficient than with a pyrimidine at the +1 position. Therefore, a purine nucleotide at the +1 position is a dominant determinant for M1GS cleavage site selection and cleavage efficiency. This observation is consistent with the notion that a guanine at the +1 position, which has been found in most bacterial pre-tRNAs, serves as the guide nucleotide for M1 RNA and RNase P cleavage in *E. coli* [24]. To meet this requirement, we have provided an option in the tool to choose either purine (A or G) at the +1 position.

#### Length of the desired GS

The optimal length of the GS can vary from 12-19 nt [23], although M1GS can cleave shorter RNA substrates [24]. The longer the length of a GS, the higher the specificity of the M1GS targeting but as the GS length increases, the turnover number of the M1GS ribozyme may be negatively impacted due to tighter M1GS binding to the target RNA. By contrast, shorter GSs may lead to an enhanced M1GS turnover number but could result in more off-target cleavages. In addition to the length of GS, the GC% of GS also contributes to the M1GS ribozyme cleavage efficiency because GC% determines the strength of base-pairing. For these reasons, the users may try GSs of different lengths if they deem necessary. Based on the length of the guide sequence chosen (excluding 3’ NCCA), target sites of an equal length in the target RNA will be predicted. Note that the length of GS does not change the positions or number of the M1GS target sites in the target RNA.

### The desirable length of the GS-binding target RNA from the -1 position to be in a single-stranded configuration

For M1GS to efficiently bind and cleave the target RNA, it is preferable to choose a region of the target RNA that is likely to exist in a single-stranded form. To identify such regions in the target RNA, past studies have used chemical methods [23, 30] and prediction software [25, 27]. The *M1GS target site finder* tool is linked to the publicly available RNAfold software tool (http://rna.tbi.univie.ac.at/cgi-bin/RNAWebSuite/RNAfold.cgi).The tool, by default, uses the RNAfold-predicted secondary structure of the target RNA, which is then used to identify regions in the target RNA that are predicted to exist in accessible conformation. The default settings of the RNAfold tool used for secondary structure prediction are shown in Figure S2. Alternatively, the users can provide RNA secondary structure input in Vienna format obtained from any other RNA secondary structure prediction tool of their choice.

To find such desirable target RNA sites, users should provide a desirable length of the GS-binding target RNA from the -1 position to be in a single-stranded configuration as an input. For example, if the length of the GS is 14 nt, one can start by providing the input specifying the desirable length of the target RNA in a single-stranded form as 14 nt. Although it is desirable to use target RNA sites that would likely exist in a single-stranded form *in vivo*, it is often not possible to find such ideal sites in target RNAs based on the *in silico* secondary structure prediction. M1GS could still bind and cleave such RNAs [23, 25]. Therefore, if no target sites are found with the desired target RNA length in a single-stranded form, the end user is recommended to keep reducing the required length until a minimum number of desired target sites is found. In cases where the software predicts that no region of the input RNA sequence that matches M1GS requirements exists in a single-stranded configuration, the input for the desirable length of a single-stranded target RNA can be put at ‘0’. The M1GS could still cleave such targets, although it might affect cleavage efficiency. The reasoning is that the secondary structures predicted by software may not reflect the exact *in vivo* scenarios. Therefore, GSs may still bind and cleave such target sites [23, 25]. Alternatively, the predicted RNA secondary structures may be weak and thus could be disrupted in the presence of the customized M1GS RNA, favoring the base-pairing between the target RNA and the M1GS. The overall approach to choosing this or the other features could be to start with the ideal combination of features and start relaxing the non-essential features as needed, depending upon the output. Alternatively, the users could also start with relaxed features first and then keep increasing the stringency until the desired M1GS target sites are found.

#### Secondary structure of the target RNA

Providing the secondary structure of the target RNA is an optional field in the tool. If this field is left empty, the tool uses the RNAFold web server by default to predict the secondary structure of the target RNA from the given input sequence (DNA or RNA). If secondary structure predictions from other software are used, they should be entered in the Vienna format in the specified input space in the tool. For example, another RNA folding prediction software, Mfold, provides the secondary structure output in multiple formats, including the Vienna format, which is suitable for this tool. Often, different prediction software give different predicted secondary structures for the same RNA sequence, as demonstrated below in the Results and Discussion sections.

#### Unpaired nucleotide at the -1 position

In consistent with the recommended unpaired RCCA or NCCA at the 3’ end of GS [15, 46-48], we have an additional feature to choose if the nucleotide at the -1 position, as per the predicted secondary structure, is base-paired or free. The purpose is to keep the cleavage site on the target RNA in a single-stranded form, as base-pairing of the nucleotide at the -1 position may act as a structural anti-determinant affecting cleavage rate and specificity [49]. However, cleavage can occur even when the nucleotide at the -1 position is paired [25], but cleavage efficiency may be low in such cases.

#### Unpaired CCA-3’

To mimic *E. coli* pre-tRNA substrates, an unpaired RCCA or NCCA is added at the 3’ end of the M1GS RNA [15, 46-48]. Similar to the nucleotide at the -1 position, keeping CCA unpaired is desirable for efficient cleavage and site selection [49]. The reason is that base-pairing of this CCA with one of the M1 RNA domains is essential for catalytic activity and site selection. This base-pairing would get impeded if 3’ CCA base-pairs with the target RNA [49]. Therefore, we recommend users to choose unpaired CCA at the 3’ end, but still provide an option in our tool to identify sites without unpaired 3’ CCA if desired by users for some experimental reasons.

#### Avoiding off-target effects

The M1GS target sites chosen to cleave a target RNA should ideally be unique to avoid off-target effects. To avoid off-target effects, the user is recommended to verify that the chosen target site(s) do not hybridize with any other RNAs in the targeted cell type, using available relevant bioinformatic tools.

#### Running the tool

Upon entering the input RNA sequence (with or without secondary structure data) and selected the desired features of the GS and the target RNA, the user can run the tool by clicking ‘find target sites’.

#### Output

The output provides a list of all possible M1GS target sites based on the provided inputs. The list consists of three components: target sites with a specific nucleotide at the -1 position (e.g., U) in the target RNA, the target site sequences (from the +1), and the GC content of the target sites, as described below with screenshots from the tool.

### Target site selection and designing guide sequence

M1GS can be employed to cleave both coding and non-coding RNA. If the mRNA of any protein-coding gene is targeted, targeting the region towards the 5’ end of the mRNA is preferred because a single cleavage in the 5’ end would likely result in the loss of the 5’ protective guanosine cap and subsequent degradation of the mRNA by ribonucleases. Instead, if the 3’ end of the mRNA is targeted, most of the target mRNA might still be available for some translation to occur, rendering gene knockdown less efficient. In such cases, some truncated proteins may form, whose effects are unpredictable. Therefore, while choosing GS for any mRNA, preference may be given to the sites proximal to the 5’ end of the mRNA. In this study, we chose to target human 28S rRNA, which is the most abundant RNA species and a non-coding RNA.

After choosing a target site, a GS should be designed, containing a sequence complementary to the target site and either RCCA or NCCA at the 3’ end [15, 46-48]. The presence of CCA at the 3’ end of GS is critical for efficient cleavage but not required for cleavage [24]. Although all three nucleotides are very critical, CC (two cytosines) are more critical than A in determining the cleavage efficiency, but they are not important for cleavage site selection [24].

### Prediction of M1GS target sites for human 28S rRNA

To demonstrate that the bioinformatic tool we designed works as expected, we used it to identify M1GS target sites in the human 28S rRNA sequence. A primary reason to choose to target 28S rRNA is to propose M1GS-mediated rRNA downregulation as a potential anticancer modality for cancers in which 45S rRNA genes are upregulated [31-33] and RNA Pol I transcription has been therapeutically targeted [34-40]. The understanding has been that reduced protein synthesis in the cancer cells would result in antiproliferation effects. Furthermore, among different rRNAs, we chose 28S because it possesses catalytic activity that forms peptide bonds between amino acids during protein synthesis.

We obtained a publicly available 28S rDNA sequence (GenBank accession number: KY962518.1) (Figure S3), converted it to the corresponding RNA sequence, and provided it as the input sequence in the tool. We started with stringent input options. For the nucleotide at the -1 position, pyrimidines (C and U) were selected, and for the nucleotide at the +1 position, purine (G) was selected. Lastly, an unpaired CCA at the 3’ end and an unpaired nucleotide at the -1 position were opted for. The output listed multiple target sites, of which one at position 2862 (Figure 3, enclosed by a red rectangle ) was used to generate a guide sequence of 15 nt for cleaving 28S rRNA. A reason to pick this target site is that it is located close to the center of the 28S rRNA sequence. The notion is that if we cleave 28S rRNA right in the middle, maximum damage to the RNA can be inflicted, and, relatively, it is easier to design suitable primers for qPCR analysis of the target region. Next, we demonstrate that the tool provides more target sites for 28S rRNA in the output if we relax some input options. For example, when we excluded the requirement of unpaired CCA at the 3’ end and an unpaired nucleotide at the -1 position, the output listed more target sites than before (compare outputs of Figure 3 with Figure S4).

**Figure 3:**
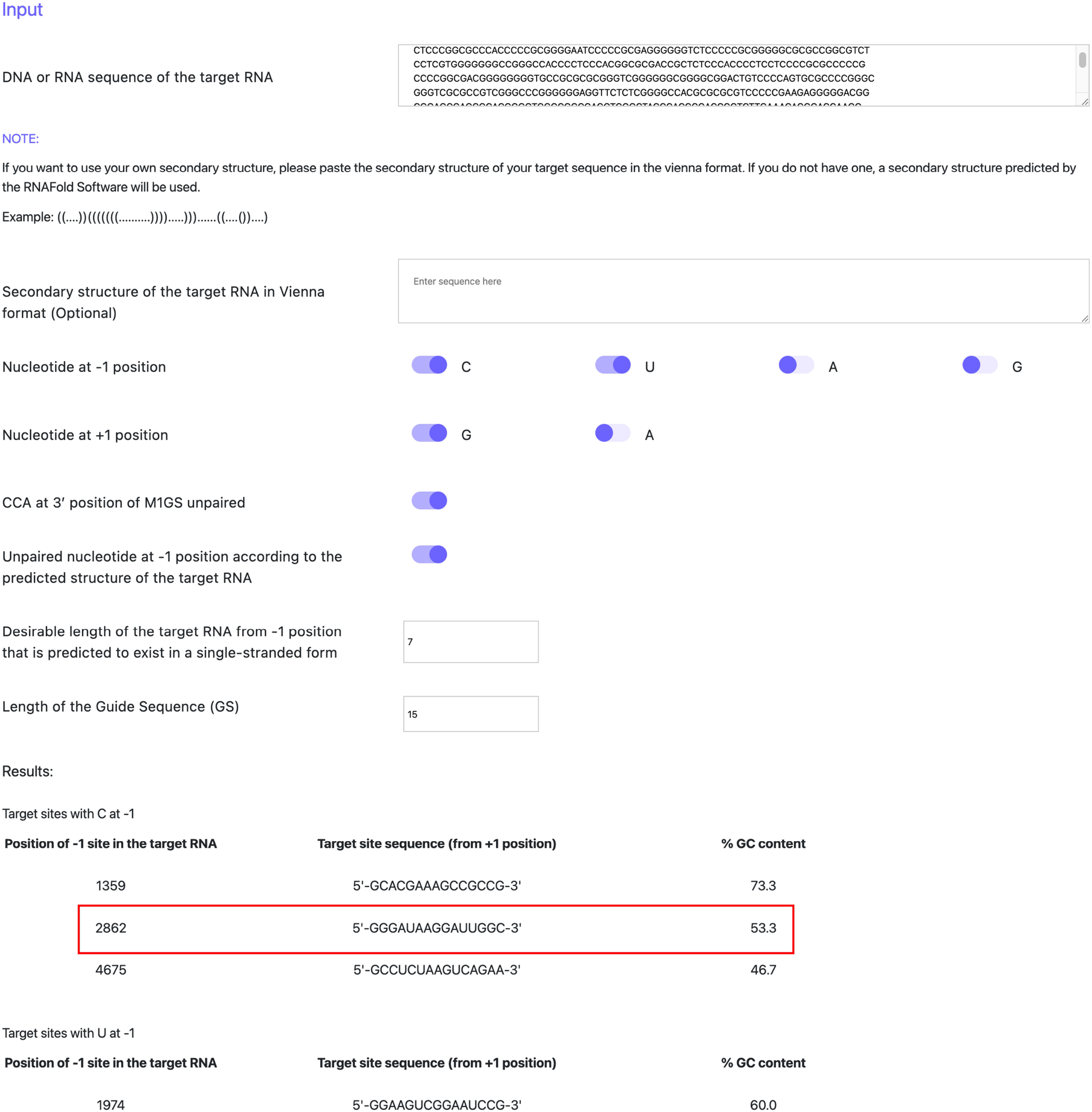
Identification of M1GS target sites for human 28S rRNA using the tool. Screenshot from the *M1GS target site finder* tool that shows input options chosen and output as a list of M1GS target sites for 28S rRNA. The input options chosen here are more stringent than the ones shown in Figure S4. Note:For better visualization, this figure was created by merging multiple cropped snapshots instead of showing a single snapshot.

### M1GS-mediated downregulation of 28S rRNA in a lung cancer cell line

The GS of 15 nt designed to target 28S rRNA was fused to wild-type M1 RNA (hereafter referred to as M1GS^28SrRNA^) or enzymatically inactive mutant M1 RNA (hereafter referred to as ΔM1GS^28SrRNA^), which served as a negative control to delineate the antisense effects, if any, that could arise due to base-pairing of the GS with the target RNA [9, 23]. A cartoon depicting different elements of the designed M1GS constructs is given in Figure S5A. M1GS can be expressed either with a 3’ overhang (Figure S5B) or without a 3’ overhang sequence by inserting a self-cleaving hammerhead ribozyme at the 3’ end of GS (Figure S5C). DNA sequences of M1GS cassettes, sequence alignment of the active and inactive M1 RNAs, and a PCR assay to differentiate the active and inactive M1 RNAs are provided in Figure S6.

Usage of a self-cleaving hammerhead ribozyme at the 3’ end of the GS has been recommended for an efficient cleavage because additional sequences downstream of the 3’ CCA sequence lower the cleavage rate of the substrates by M1 RNA [23]. We introduced a self-cleaving hammerhead (HH) ribozyme [27], downstream of the M1GS, to ensure a free 3′ end for efficient cleavage of the 28S rRNA by the M1GS^28SrRNA^. We transfected the human lung cancer cell line A549 with M1GS^28SrRNA^ and ΔM1GS^28SrRNA^ in two biological replicates. Following transfection, DNA-free RNA was obtained and used to perform quantitative PCR (qPCR) and semiquantitative RT-PCR assays for 28S rRNA, M1GS RNAs, and internal controls. Non-transfected cells were used as controls.

The qPCR assay results showed a significant reduction in the levels of 28S rRNA in M1GS^28SrRNA^-expressing cells compared to ΔM1GS^28SrRNA^-expressing and the non-transfected cells (Figure 4A). In both biological replicates, we found 28S rRNA downregulation, but in one replicate we observed greater 28S rRNA reduction than the other. We used the averages of the two biological replicates for a box plot analysis (Figure 4A). To further confirm the results, we carried out a semiquantitative RT-PCR assay using cDNA from the biological replicate which showed a greater reduction of 28S rRNA (Figure 4B). We observed similar reduction of 28S rRNA, not GAPDH (an internal control), in the transfected cells expressing the active M1GS^28SrRNA^ compared to the cells expressing ΔM1GS^28SrRNA^ or the non-transfected cells (Figure 4B, image panels 1-4). We confirmed successful transfection of the cells through DNA PCR amplification of both M1GSs and GAPDH (as a control) (Figure 4B, image panels 5-6), and a separate RT-PCR assay to confirm the expression of both M1GSs (Figure 4B, image panels 7-8 & 9-10), and GAPDH controls (Figure 4B, image panels 13-14). Further, we demonstrated self-cleavage of the HH ribozyme that was part of the M1GS constructs (Figure 4B: For M1GS^28SrRNA^, compare gel panel 7, lane 3 with panel 11, lane 3. For ΔM1GS^28SrRNA^, compare gel panel 9, lane 2 with panel 11, lane 2). We also assayed for M1GS expression using qPCR (Figure S7). Both RT-PCR and qPCR data (Figure 4B and Figure S7) revealed that ΔM1GS^28SrRNA^ is expressed at higher levels than the ΔM1GS^28SrRNA^ in the transfected cells. It appears that, with transient transfections, the expression levels of the constructs are inconsistent, which also explains the differential downregulation of 28S rRNA between the two biological replicates. So, these results indicate that higher reductions of 28S rRNA are likely possible with increased expression levels of M1GS^28SrRNA^.

**Figure 4:**
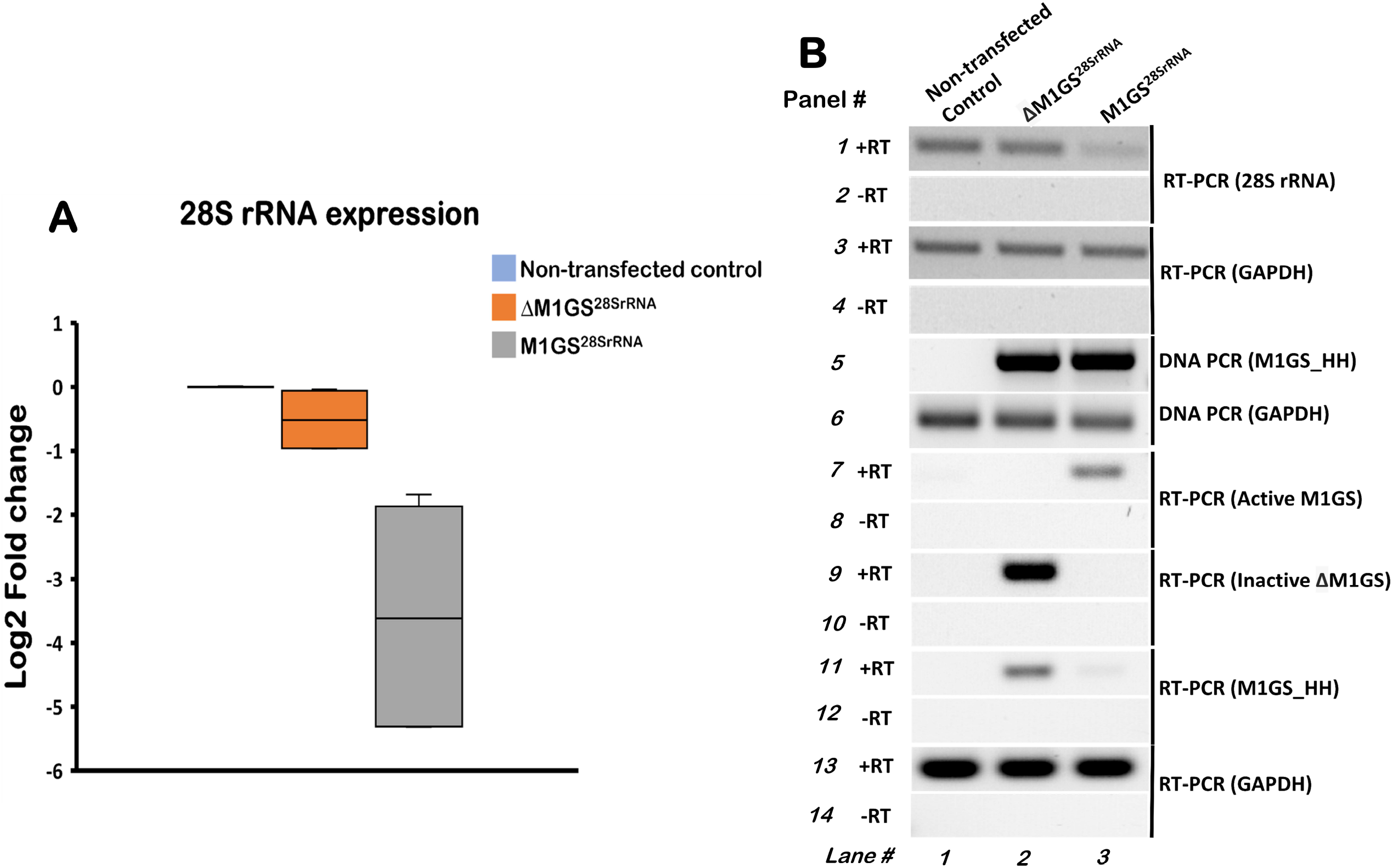
M1GS-mediated downregulation of 28S rRNA in human lung cancer cell line. (A) A box plot analysis of qPCR data depicts relative 28S rRNA levels in human A549 cell line expressing different M1GS constructs and non-transfected controls. 28S rRNA expression levels are normalized to those of non-transfected controls. (B) Gel images show semiquantitative RT-PCR analysis of 28S rRNA and GAPDH, PCR analysis of M1GS constructs and GAPDH, RT-PCR analysis of M1GS, GAPDH, and the hammerhead ribozyme. Primers specific to active and inactive M1GS were used for their amplification.

### Further validation of the *M1GS target site finder* tool

We tested our tool to validate target sites for two previously published target RNAs that were cleaved by customized M1GSs. One of the targets was the thymidine kinase (TK) mRNA of the human herpes simplex virus 1 (HSV-1) [23]. The gene sequence and a screenshot of the output showing a list of M1GS targets sites identified by the tool are provided in Figure S8, in which the site enclosed in a red rectangle was used in the study [23].

The other target we tested is the 5’ UTR of the hepatitis C virus (HCV) [25]. For this target RNA, we used both RNAfold and Mfold software to predict the secondary structure of the HCV RNA. The results demonstrate that different software provide different secondary structure predictions for the same RNA sequence (Figure S9). For example, for a 13 nt GS, when Mfold or RNAfold-predicted secondary structure was fed to the tool with a requirement of ≥ 7 nt in a single-stranded form, the tool did not find any target sites. With a requirement of 6 nt to be in the single-stranded form, when the secondary structure as predicted by the Mfold software was fed to the tool, it provided two target sites, including the one that was used in the previous study to design a GS to cleave the target RNA (Figure S9B) [25]. However, for the same input options, when the secondary structure predicted by RNAfold was fed to the tool, no M1GS target sites were identified by the tool because, according to the RNAfold prediction, target sites with a stretch of 6 nt or less (all the way to 1 nt) do not exist in a single-stranded form (Figure S9C). However, for the same RNAfold-predicted secondary structure, when the input of single-stranded form requirement of the target RNA was set to ‘0’, the tool predicted three M1GS target sites, including the one used in the study (Figure S9D) [25].

## Discussion

We describe here a user-friendly Python script-based bioinformatic tool we designed to identify M1GS cleavage sites in any target RNA. Overall, the tool offers various options for selecting GS features as per the recommended parameters to optimally meet M1GS requirements and users’ choices to experiment with. The tool has been made publicly available for use and a link to access it is provided below.

Human rRNA undergoes more than 200 modifications, of which some are reported to be altered in multiple cancers. Examples include pseudouridylation and 2’-O-methylation [50-52]. It remains to be tested if any of these or other rRNA modifications near the cleavage site in the target RNA have any impact on M1GS cleavage. Such studies would enable the incorporation of more features into the bioinformatic tool described here. For example, if any of the rRNA modifications negatively impact M1GS cleavage, the tool can be linked to available databases of rRNA modifications in both cancerous and non-cancerous cells to avoid target regions around sites carrying potentially impeding rRNA modifications. Similarly, the tool can be further linked to the human transcriptome of specific cell types to avoid potential off-targets. We have deposited the script on GitHub (https://github.com/navinfreeze/m1gstargetfinder), which we permit for unrestricted download and modification as deemed necessary by users.

Customized ribozymes have seldom been used as gene-targeting tool or antiviral agents, despite their versatile utility [25, 53, 54]. We hope that the availability of a user-friendly bioinformatic tool to identify M1GS target sites would encourage more users to employ the M1GS tool in RNA-cleaving/reverse genetic studies and to further modify the tool as more data emerge on M1GS functionality.

Of note, M1GS has not been previously used to downregulate RNA as abundant as rRNA, which constitutes ∼ 80% of total RNA in human cells [55]. Therefore, the efficient downregulation of abundant 28S rRNA observed in this study suggests a high turnover number for the M1GS ribozyme. In comparison, the available RNA-cleaving tools, RNAi and Cas13a, have a modest turnover number of ∼10 [56, 57]. Therefore, our findings hold a promise in developing the M1GS as the most potent RNA-cleaving tool. Further, compared with the RNA interference (RNAi) and other gene-targeting strategies, M1GS ribozymes offer several advantages. One is that they cleave the target RNA without the need to introduce any additional protein cofactors [19, 58, 59]. On the other hand, RNAi, for example, requires several cellular factors, such as Exportin V, Dicer, and Argonaute proteins, for small interfering RNA (siRNA)-mediated gene silencing [60].

Furthermore, several viruses have evolved counter-defense mechanisms to block RNAi-mediated host defense responses [61, 62]. By contrast, viruses lack counter-defense mechanisms against engineered ribozymes as they don’t encounter them naturally during infections. Moreover, modulation of ribozyme expression levels is simpler than that of the other RNA-cleaving tools, as it requires expression of only one component. On the other hand, RNAi and CRISPR-Cas-based systems require simultaneous modulation of multiple components.

Further experiments are needed to determine whether M1GS-mediated downregulation of rRNA exhibits anti-cancer activity in cancers where rRNA is upregulated. Direct downregulation of rRNA, if proven successful in exhibiting anticancer activity, may serve as a better viable alternative to targeting RNA pol I, because RNA pol I transcription may be necessary for other cellular functions [63].

The next concern is the *in vivo* delivery of M1GS constructs. Notably, previous studies have demonstrated the feasibility of systemic delivery of M1GS-containing plasmids in virus-infected mouse models using attenuated bacteria, and successfully cleaved the target viral RNA, resulting in reduced viral infections [64, 65].

Lastly, a question of interest has been, how does M1GS of bacterial origin work so efficiently in human cells? Is any human protein(s) involved in aiding M1GS cleavage of the target RNA? These interesting questions remain to be answered.

## Methods

### Prediction of the target RNA secondary structure

If the secondary structure of the target RNA is not provided by the user in the Vienna format in the form of “.”, “(“, “)”) as input, the tool uses secondary structure predictions provided from the RNAfold webserver (http://rna.tbi.univie.ac.at/cgibin/RNAWebSuite/RNAfold.cgi) [66]. The default RNAfold settings are used to predict secondary structure, and these settings are listed in Figure S2.

#### The web interface

The web interface for the tool was developed using Python3 and Django 3.1.6. The web interface mainly consists of two sections: the inputs and the output. To find the target sites, the interface runs the program in the background, and the time required for this task varies depending on the size of the target RNA. Once this task is over, the tool loads the output above ‘the inputs’ section. The M1GS target site finder tool can be accessed at the following link http://m1gstargetfinder.bits-hyderabad.ac.in:8000/m1gs/.

#### Plasmid construction

The M1GS expression cassettes designed to target human 28S rRNA consisted of H1 promoter, followed by wildtype (active) or mutant (inactive) M1 RNA, a linker region, a 15 bp guide sequence (GS) (5’-GCCAATCCTTATCCC-3’), and a self-cleaving hammerhead ribozyme (HH) (Fig. S5). The GS was designed to cleave 28S rRNA at between nts 2862 and 2863, a target site identified by the bioinformatic tool. These expression cassettes were commercially synthesised (ABclonal), and then cloned into a pRS retroviral vector using EcoRI and XhoI restriction sites. Plasmid DNA from the positive clones was isolated using *QIAprep Spin Miniprep Kit* (Qiagen), and the presence of the M1GS constructs was verified by PCR amplification of the active and inactive M1 DNA using specific primers.

#### Human cell culture

The human lung cancer cell line A549 (ATCC CCL-185) was purchased from the American Type Culture Collection (Manassas, VA, USA). The cells were cultured in Dulbecco’s Modified Eagle Medium (DMEM) (HiMedia), supplemented with 10% (v/v) fetal bovine serum (HiMedia) and 1% (w/v) penicillin streptomycin (HiMedia), at 37° in a humidified incubator with 5% CO_2_. The medium was changed every 2 days until the cells reached 70–80% confluence. For detaching the cells during passages, 0.25% (w/v) Trypsin-EDTA (HiMedia) was used.

#### Transfection

Transient transfection of A549 cells was performed using JetPRIME® kit (PolyPlus). Once the cells in a 6-well plate reached 70-80% confluency, a transfection master mix consisting of 2 μg plasmid DNA containing active (pRS-M1GS^28SrRNA^) or inactive M1GS (pRS-ΔM1GS^28SrRNA^) and 4 μl of JetPrime reagent (1:2 ratio) in 200 μL of JetPrime® buffer (PolyPlus) was added for each well and then the cells were incubated at 37° for 24 h. For transfection, the cells were seeded at a density of 2,50,000 cells per well in a 6-well plate. After transfection, cells were supplemented with additional media. The cells were harvested 24 h post-transfection and processed immediately for RNA and DNA isolation.

### DNA and RNA isolation and semiquantitative reverse transcription (RT)-PCR

Total DNA and RNA were isolated from the control and transfected cell lines using the *AllPrep DNA/ RNA/Protein Mini Kit* (Qiagen). To confirm the presence of the M1GS constructs in the transfected cells, PCR amplification of M1GS constructs and the endogenous glyceraldehyde-3-phosphate dehydrogenase (GAPDH) gene (as an internal control) was carried using initial denaturation for 2 min at 95°, followed by 30 cycles of 30 sec at 95°, 30 sec at 58°, and 30 sec at 72°, and a final extension for 3 min at 72°, in the presence of *GoTaq® Green Master Mix* (Promega).

The isolated total RNA was treated with *Turbo DNA-free* kit (Invitrogen) to eliminate any contaminating genomic DNA. One μg of DNA-free RNA was reverse transcribed to cDNA using *SuperScript III First Strand cDNA synthesis* kit (Invitrogen). One μl (cDNA from ∼50 ng DNA-free RNA) of reverse transcribed product was then used for PCR amplification of the active and the inactive M1GS ribozymes, the HH ribozyme, and GAPDH control with initial denaturation for 2 min at 95°, followed by 25 cycles of 30 sec at 95 °, 30 sec at 58 °, and 30 sec at 72 °, and a final extension for 3 min at 72 °. The same PCR profile was used for amplification of 28S rRNA, but for 23 cycles. For all semiquantitative RT-PCR reactions, RT-negative controls were included.

### Quantitative PCR (pPCR) assays

The qPCR assays were performed using a *Step One Plus* real-time PCR system (Applied Biosystems) in the presence of *TB Green® Premix Ex Taq™* (Tli RNase H Plus) (Takara). The qPCR amplification was performed using an initial denaturation at 95 ° for 10 sec, followed by 40 cycles of 15 sec at 95 ° and 60 sec at 60 °. The relative gene expression levels of 28S RNA and M1GS were calculated using the 2^(-△△CT) method (Livak & Schmittgen, 2001) by normalizing their expression levels to the expression levels of GAPDH as a reference gene. qPCR was performed in duplicates for each sample. Primer sequences used to amplify 28S rRNA span the M1GS cleavage site (2862). All the primers used in this study are listed in Figure S10.

### Statistical analysis

Data from two technical replicates from two biological replicates were used for all the calculations and data analysis. The qPCR data were used to generate a boxplot. The standard two-tailed t-test was used to analyze significant differences in the gene expression levels between the control (non-transfected) and the transfected cells expressing active or inactive M1GS. A *p* of < 0.05 was considered statistically significant.

### Web tools for sequence alignments

Alignment of the active and the inactive M1 RNA sequences was carried out using the nucleotide BLAST (blastn) tool from NCBI (https://blast.ncbi.nlm.nih.gov/Blast.cgi?PROGRAM=blastn&BLAST_SPEC=GeoBlast&PAGE_TYPE=BlastSearch).

### Other software used

Figures were made using the free version of Figma software and the licensed version of PowerPoint.

## Acknowledgements

We profusely thank Dr. Venkat Gopalan and Dr. Lien Lai, The Ohio State University (USA), for their valuable support in sharing reagents, expertise, and critique of the manuscript. We are also thankful to the CCIT division and the department of Biological Sciences and BITS-Pilani Hyderabad campus, for their support.

## Author Contributions

N.P., N.V.P, H.P., S.B., and G.M. designed research, N.P., N.V.P., and H.P. carried out research, N.P., N.V.P., H.P., and G.M. analyzed data, N.P., N.V.P., H.P., and G.M. wrote the manuscript. All authors reviewed the manuscript.

