## Supplementary Figures for "Engineered bacterial M1GS ribozyme efficiently cleaves the most abundant ribosomal RNA in a human cancer cell line"

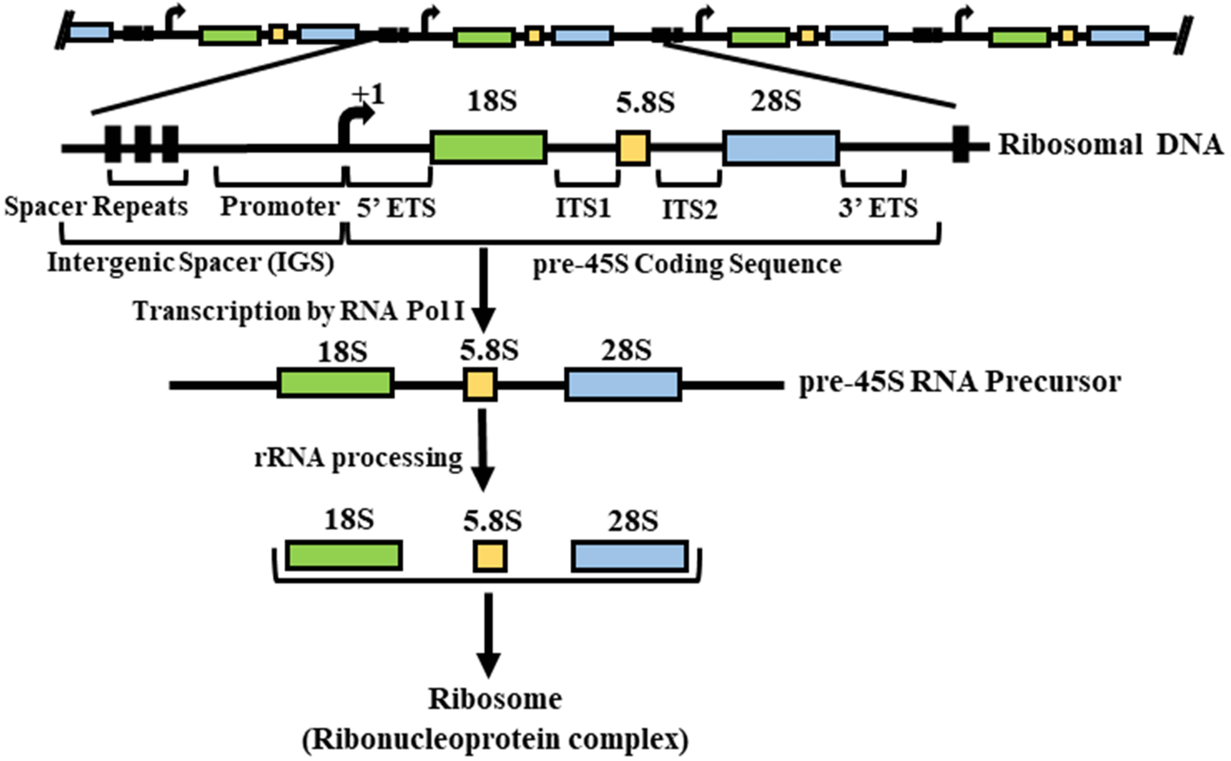


**Figure S1: Organization, transcription, and processing of human 45S rRNA genes.** The diagram shows human 45S rRNA gene organization, components of a typical human 45S rRNA gene, including the structural rRNAs, which, upon transcription and processing, become components of functional ribosomes.

Fold algorithms and basic options:

- Minimum free energy (MFE) and partition function
- Avoid isolated base pairs

Advanced options:

- Dangling end options: dangling energies on both sides of a helix in any case
- Energy Parameters: RNA parameters (Turner model, 2004)
- SHAPE reactivity data: None
- Rescale energy parameters to a given temperature (C) - 37

**Figure S2: Default settings of RNAFold tool used for secondary structure prediction**

**Target RNA: 28S ribosomal RNA gene**

Target RNA used in the paper: 5'-GGGAUAAGGAUUGGC-3'

>KY962518.1:7921-12971 Homo sapiens 28S ribosomal RNA gene

```
CGCGACCTCAGATCAGACGTGGCGACCCGCTGAATTTAAGCATATTAGTCAGCGGAGGAAAAGAAACTAACCAGGATTCCCTCAGT
AACGGCGAGTGAACAGGGAAGAGCCCAGCGCCGAATCCCCGCCCCGCGGCGGGGCGCGGGACATGTGGCGTACGGAAGACCCGCTC
CCCGGCGCCGCTCGTGGGGGGCCCAAGTCCTTCTGATCGAGGCCCAGCCCGTGGACGGTGTGAGGCCGGTAGCGGCCCCCGGCGCG
CCGGGCCCGGGTCTTCCCGGAGTCGGGTTGCTTGGGAATGCAGCCCAAAGCGGGTGGTAAACTCCATCTAAGGCTAAATACCGGCA
CGAGACCGATAGTCAACAAGTACCGTAAGGGAAAGTTGAAAAGAACTTTGAAGAGAGAGTTCAAGAGGGCGTGAAACCGTTAAGAG
TAAACGGGTGGGGTCCGCGCAGTCCGCCCGGAGGATTCAACCCGGCGGCGGGTCCGGCCGTGTCGGCGGCCCGGCGGATCTTTCCC
GCCCCCCGTTCCTCCCGACCCCTCCACCCGCCCTCCCTTCCCCCGCCGCCCCTCCTCCTCCCCGGAGGGGGCGGGCTCCGGCG
GGTGCGGGGGTGGGCGGGCGGGGCCGGGGGTGGGGTCGGCGGGGGACCGTCCCCCGACCGGCGACCGGCCGCCGCCGGGCGCATTT
CCACCGCGGCGGTGCGCCGCGACCGGCTCCGGGACGGCTGGGAAGGCCCGGCGGGGAAGGTGGCTCGGGGGGCCCCGTCCGTCCGT
CCGTCCGTCCTCCTCCTCCCCCGTCTCCGCCCCCCGGCCCCGCGTCCTCCCTCGGGAGGGCGCGCGGGTCGGGGCGGCGGCGGCGG
TGGCGGCGGCGGCGGCGGCGGCGGGACCGAAACCCCCCCGAGTGTTACAGCCCCCCCGGCAGCAGCACTCGCCGAATCCCGGGGCG
AGGGAGCGAGACCCGTCGCCGCGCTCTCCCCCCTCCCGGCGCCCACCCCCGCGGGGAATCCCCCGCGAGGGGGGTCTCCCCCGCGG
GGGCGCGCCGGCGTCTCCTCGTGGGGGGGCCGGGCCACCCCTCCCACGGCGCGACCGCTCTCCCACCCCTCCTCCCCGCGCCCCCG
CCCCGGCGACGGGGGGGGTGCCGCGCGCGGGTCGGGGGGCGGGGCGGACTGTCCCCAGTGCGCCCCGGGCGGGTCGCGCCGTCGGG
CCCGGGGGGAGGTTCTCTCGGGGCCACGCGCGCGTCCCCCGAAGAGGGGGACGGCGGAGCGAGCGCACGGGGTCGGCGGCGACGTC
GGCTACCCACCCGACCCGTCTTGAAACACGGACCAAGGAGTCTAACACGTGCGCGAGTCGGGGGCTCGCACGAAAGCCGCCGTGGC
GCAATGAAGGTGAAGGCCGGCGCGCTCGCCGGCCGAGGTGGGATCCCGAGGCCTCTCCAGTCCGCCGAGGGCGCACCACCGGCCCG
TCTCGCCCGCCGCGCCGGGGAGGTGGAGCACGAGCGCACGTGTTAGGACCCGAAAGATGGTGAACTATGCCTGGGCAGGGCGAAGC
CAGAGGAAACTCTGGTGGAGGTCCGTAGCGGTCCTGACGTGCAAATCGGTCGTCCGACCTGGGTATAGGGGCGAAAGACTAATCGA
ACCATCTAGTAGCTGGTTCCCTCCGAAGTTTCCCTCAGGATAGCTGGCGCTCTCGCAGACCCGACGCACCCCCGCCACGCAGTTTT
ATCCGGTAAAGCGAATGATTAGAGGTCTTGGGGCCGAAACGATCTCAACCTATTCTCAAACTTTAAATGGGTAAGAAGCCCGGCTC
GCTGGCGTGGAGCCGGGCGTGGAATGCGAGTGCCTAGTGGGCCACTTTTGGTAAGCAGAACTGGCGCTGCGGGATGAACCGAACGC
CGGGTTAAGGCGCCCGATGCCGACGCTCATCAGACCCCAGAAAAGGTGTTGGTTGATATAGACAGCAGGACGGTGGCCATGGAAGT
CGGAATCCGCTAAGGAGTGTGTAACAACTCACCTGCCGAATCAACTAGCCCTGAAAATGGATGGCGCTGGAGCGTCGGGCCCATAC
CCGGCCGTCGCCGGCAGTCGAGAGTGGACGGGAGCGGCGGGGGCGGCGCGCGCGCGCGCGTGTGGTGTGCGTCGGAGGGCGGCG
GCGGCGGCGGGGGTGTGTGGGGTCCTCCCCCGCCCCCCCCCACGCCTCCTCCCCTCCTCCCGCCCACGCCCCGCTCCCCGCCCCCG
GAGCCCCGCGGACGCTACGCCGCGACGAGTAGGAGGGCCGCTGCGGTGAGCCTTGAAGCCTAGGGCGCGGGCCCGGGTGGAGCCGC
CGCAGGTGCAGATCTTGGTGGTAGTAGCAAATATTCAAACGAGAACTTTGAAGGCCGAAGTGGAGAAGGGTTCCATGTGAACAGCA
GTTGAACATGGGTCAGTCGGTCCTGAGAGATGGGCGAGCGCCGTTCTGAAGGGACGGGCGATGGCCTCCGTTGCCCTCGGCCGATC
GAAAGGGAGTCGGGTTCAGATCCCCGAATCCGGAGTGGCGGAGATGGGCGCCGCGAGGCGTCCAGTGCGGTAACGCGACCGATCCC
GGAGAAGCCGGCGGGAGCCCCGGGGAGAGTTCTCTTTTCTTTGTGAAGGGCAGGGCGCCCTGGAATGGGTTCGCCCCGAGAGAGGG
GCCCGTGCCTTGGAAAGCGTCGCGGTTCCGGCGGCGTCCGGTGAGCTCTCGCTGGCCCTTGAAAATCCGGGGGAGAGGGTGTAAAT
CTCGCGCCGGGCCGTACCCATATCCGCAGCAGGTCTCCAAGGTGAACAGCCTCTGGCATGTTGGAACAATGTAGGTAAGGGAAGTC
GGCAAGCCGGATCCGTAACTTCGGGATAAGGATTGGCTCTAAGGGCTGGGTCGGTCGGGCTGGGGCGCGAAGCGGGGCTGGGCGCG
CGCCGCGGCTGGACGAGGCGCCGCCGCCCCCCCCACGCCCGGGGCACCCCCCTCGCGGCCCTCCCCCGCCCCACCCCGCGCGCGCC
GCTCGCTCCCTCCCCACCCCGCGCCCTCTCTCTCTCTCTCTCCCCCGCTCCCCGTCCTCCCCCCTCCCCGGGGGAGCGCCGCGTGG
GGGCGGCGGCGGGGGGAGAAGGGTCGGGGCGGCAGGGGCCGGCGGCGGCCGCCGCGGGGCCCCGGCGGCGGGGGCACGGTCCCCCG
CGAGGGGGGCCCGGGCACCCGGGGGGCCGGCGGCGGCGGCGACTCTGGACGCGAGCCGGGCCCTTCCCGTGGATCGCCCCAGCTGC
GGCGGGCGTCGCGGCCGCCCCCGGGGAGCCCGGCGGGCGCCGGCGCGCCCCCCCCCCACCCCACGTCTCGTCGCGCGCGCGTCCGC
TGGGGGCGGGGAGCGGTCGGGCGGCGGCGGTCGGCGGGCGGCGGGGCGGGGCGGTTCGTCCCCCCGCCCTACCCCCCCGGCCCCGT
CCGCCCCCCGTTCCCCCCTCCTCCTCGGCGCGCGGCGGCGGCGGCGGGCGGCGGAGGGGCCGCGGGCCGGTCCCCCCCGCCGGGTC
CGCCCCCGGGGCCGCGGTTCCGCGCGGCGCCTCGCCTCGGCCGGCGCCTAGCAGCCGACTTAGAACTGGTGCGGACCAGGGGAATC
CGACTGTTTAATTAAAACAAAGCATCGCGAAGGCCCGCGGCGGGTGTTGACGCGATGTGATTTCTGCCCAGTGCTCTGAATGTCAA
AGTGAAGAAATTCAATGAAGCGCGGGTAAACGGCGGGAGTAACTATGACTCTCTTAAGGTAGCCAAATGCCTCGTCATCTAATTAG
TGACGCGCATGAATGGATGAACGAGATTCCCACTGTCCCTACCTACTATCCAGCGAAACCACAGCCAAGGAACGGGCTTGGCGGA
ATCAGCGGGGAAAGAAGACCCTGTTGAGCTTGACTCTAGTCTGGCACGGTGAAGAGACATGAGAGGTGTAGAATAAGTGGGAGGCC
CCCGGCGCCCCCCCGGTGTCCCCGCGAGGGGCCCGGGGCGGGGTCCGCCGGCCCTGCGGGCCGCCGGTGAAATACCACTACTCTGA
TCGTTTTTTCACTGACCCGGTGAGGCGGGGGGGCGAGCCCCGAGGGGCTCTCGCTTCTGGCGCCAAGCGCCCGGCCGCGCGCCGGC
CGGGCGCGACCCGCTCCGGGGACAGTGCCAGGTGGGGAGTTTGACTGGGGCGGTACACCTGTCAAACGGTAACGCAGGTGTCCTAA
GGCGAGCTCAGGGAGGACAGAAACCTCCCGTGGAGCAGAAGGGCAAAAGCTCGCTTGATCTTGATTTTCAGTACGAATACAGACCG
TGAAAGCGGGGCCTCACGATCCTTCTGACCTTTTGGGTTTTAAGCAGGAGGTGTCAGAAAAGTTACCACAGGGATAACTGGCTTGT
GGCGGCCAAGCGTTCATAGCGACGTCGCTTTTTGATCCTTCGATGTCGGCTCTTCCTATCATTGTGAAGCAGAATTCACCAAGCGT
TGGATTGTTCACCCACTAATAGGGAACGTGAGCTGGGTTTAGACCGTCGTGAGACAGGTTAGTTTTACCCTACTGATGATGTGTTG
TTGCCATGGTAATCCTGCTCAGTACGAGAGGAACCGCAGGTTCAGACATTTGGTGTATGTGCTTGGCTGAGGAGCCAATGGGGCGA
AGCTACCATCTGTGGGATTATGACTGAACGCCTCTAAGTCAGAATCCCGCCCAGGCGGAACGATACGGCAGCGCCGCGGAGCCTCG
GTTGGCCTCGGATAGCCGGTCCCCCGCCTGTCCCCGCCGGCGGGCCGCCCCCCCCTCCACGCGCCCCGCGCGCGCGGGAGGGCGCG
TGCCCCGCCGCGCGCCGGGACCGGGGTCCGGTGCGGAGTGCCCTTCGTCCTGGGAAACGGGGCGCGGCTGGAAAGGCGGCCGCCCC
CTCGCCCGTCACGCACCGCACGTTCGTGGGGAACCTGGCGCTAAACCATTCGTAGACGACCTGCTTCTGGGTCGGGGTTTCGTACG
TAGCAGAGCAGCTCCCTCGCTGCGATCTATTGAAAGTCAGCCCTCGACACAAGGGTTTGTC
```

**Figure S3: Human 28S rRNA sequence.** The sequence highlighted in yellow is the target site for M1GS binding

Nucleotide at -1 position | C | U | A | G

Nucleotide at +1 position | G | A

CCA at 3′ position of M1GS unpaired

Unpaired nucleotide at -1 position according to the predicted structure of the target RNA

Desirable length of the target RNA from -1 position that is predicted to exist in a single-stranded form: 7

Length of the Guide Sequence (GS): 15

Results:

Target sites with C at -1

| Position of -1 site in the target RNA | Target site sequence (from +1 position) | % GC content |
|---|---|---|
| 1359 | 5'-GCACGAAAGCCGCCG-3' | 73.3 |
| 2691 | 5'-GGUUCCGGCGGCGUC-3' | 80.0 |
| 2854 | 5'-GUAACUUCGGGAUAA-3' | 40.0 |
| 2862 | 5'-GGGAUAAGGAUUGGC-3' | 53.3 |
| 3840 | 5'-GAAACCACAGCCAAG-3' | 53.3 |
| 4675 | 5'-GCCUCUAAGUCAGAA-3' | 46.7 |
| 4943 | 5'-GCUAAACCAUUCGUA-3' | 40.0 |

Target sites with U at -1

| Position of -1 site in the target RNA | Target site sequence (from +1 position) | % GC content |
|---|---|---|
| 1314 | 5'-GAAACACGGACCAAG-3' | 53.3 |
| 1974 | 5'-GGAAGUCGGAAUCCG-3' | 60.0 |
| 1998 | 5'-GUGUAACAACUCACC-3' | 46.7 |
| 3939 | 5'-GUAGAAUAAGUGGGA-3' | 40.0 |
| 4272 | 5'-GAUCUUGAUUUUCAG-3' | 33.3 |
| 4421 | 5'-GAUCCUUCGAUGUCG-3' | 53.3 |
| 4671 | 5'-GAACGCCUCUAAGUC-3' | 53.3 |

**Figure S4: Identification of M1GS target sites for human 28S rRNA using the tool.**
Screenshots from the *M1GS target site finder* tool that show lists of M1GS target sites for 28S rRNA. Shows results for less stringent input options. The input options chosen here are less stringent than the ones shown in Figure 3.
Note: For better visualization, this figure was created by merging multiple cropped snapshots instead of showing a single snapshot.

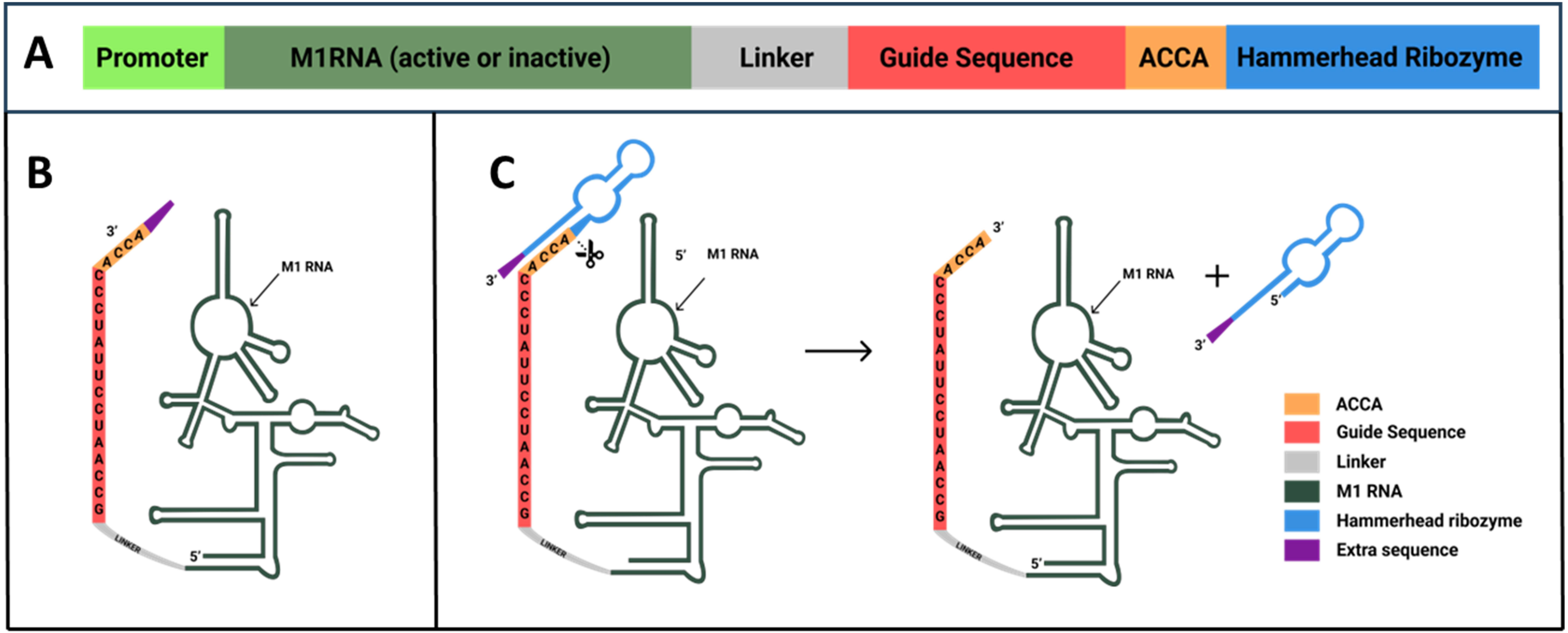


**Figure S5**: **M1GS cleavage strategy and its structural components:** (A) The cartoon depicts different components of the M1GS constructs used in this study. M1GS cleavage strategy (B) with a 3′ overhang and (C) without 3′ overhang, accomplished by introducing a self-cleaving hammerhead ribozyme at the 3′ end.

**A** **H1TetO2 promoter (human H1 RNA)**

GAATTCAATATTTGCATGTCGCTATGTGTTCTGGGAAATCACCATAAACGTGAAATCCCTATCAGTGA
TAGAGACTTATAAGTTCCCTATCAGTGATAGAGATCC

**B** **Linker sequence**

AAGCTTGGCGTAATCATGGTCATA- 24 bp

**C** **Hammerhead ribozyme sequence**

GTCGACATCTGAAACCCGATGTCCTGATGAGCTCAAGGTAGAGCGAAACTGGTCTCGAG

**D** **Sequence alignment of active and inactive M1 RNA**

```
Active     1    GAAGCTGACCAGACAGTCGCCGCTTCGTCGTCGTCCTCTTCGGGGGAGACGGGCGGAGGG   60
                |||||||||||||||||||||||||| |||||||||||||||| |||||||| |||||||
Inactive   1    GAAGCTGACCAGACAGTCGCCGCTTCATCGTCGTCCTCTTCGGAGGAGACGGACGGAGGG   60

Active     61   GAGGAAAGTCCGGGCTCCATAGGGCAGGGTGCCAGGTAACGCCTGGGGGGGAAACCCACG   120
                |||||||||||||||| |||||||||||||||||||||||||||||||||||||||||||
Inactive   61   GAGGAAAGTCCGGGCTTCATAGGGCAGGGTGCCAGGTAACGCCTGGGGGGGAAACCCACG   120

Active     121  ACCAGTGCAACAGAGAGCAAACCGCCGATGGCCCGCGCAAGCGGGATCAGGTAAGGGTGA   180
                |||||||||||||||||||||||||||||||||||||||||||||||| || ||||||||
Inactive   121  ACCAGTGCAACAGAGAGCAAACCGCCGATGGCCCGCGCAAGCGGGATCGGGCAAGGGTGA   180

Active     181  AAGGGTGCGGTAAGAGCGCACCGCGCGGCTGGTAACAGTCCGTGGCACGGTAAACTCCAC   240
                ||||||||||||||||||||||||||||||||||||||||||||||||||||||||||||
Inactive   181  AAGGGTGCGGTAAGAGCGCACCGCGCGGCTGGTAACAGTCCGTGGCACGGTAAACTCCAC   240

Active     241  CCGGAGCAAGGCCAAATAGGGGTTCATAAGGTACGGCCCGTACTGAACCCGGGTAGGCTG   300
                ||||||||||||||||||||||||||||||||||||||||||||||||||||||||||||
Inactive   241  CCGGAGCAAGGCCAAATAGGGGTTCATAAGGTACGGCCCGTACTGAACCCGGGTAGGCTG   300
                                                              <.............
Active     301  CTTGAGCCAGTGAGCGATTGCTGGCCTAGATGAATGACTGTCCACGACAGAACCCGGCTT   360
                |||| |||||||||||||||||||||||||||||||||||||||  ||      |  |
Inactive   301  CTTGGGCCAGTGAGCGATTGCTGGCCTAGATGAATGACTGTCCACGCTAGCGAGATGTAT   360
                                                              <-------------
                ................
Active     361  ATCGGTCAGTTTCACCT   377
                ||||||||||||||||
Inactive   361  CTCGGTCAGTTTCACCT   377
                ----------------
```

**E**

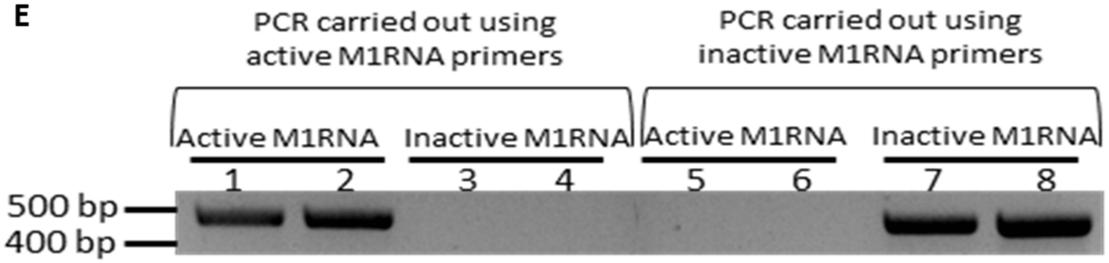


**Figure S6: Sequences of different components of M1GS constructs.** (A) H1TetO2 promoter (B) Linker sequence (C) Hammerhead ribozyme (D) An alignment of the active vs inactive M1 sequences. The arrows depict the reverse primer binding sites used to differentiate the active from the inactive M1 RNA. (E) An agarose gel image shows unique PCR amplification of the active and the inactive M1 RNA.

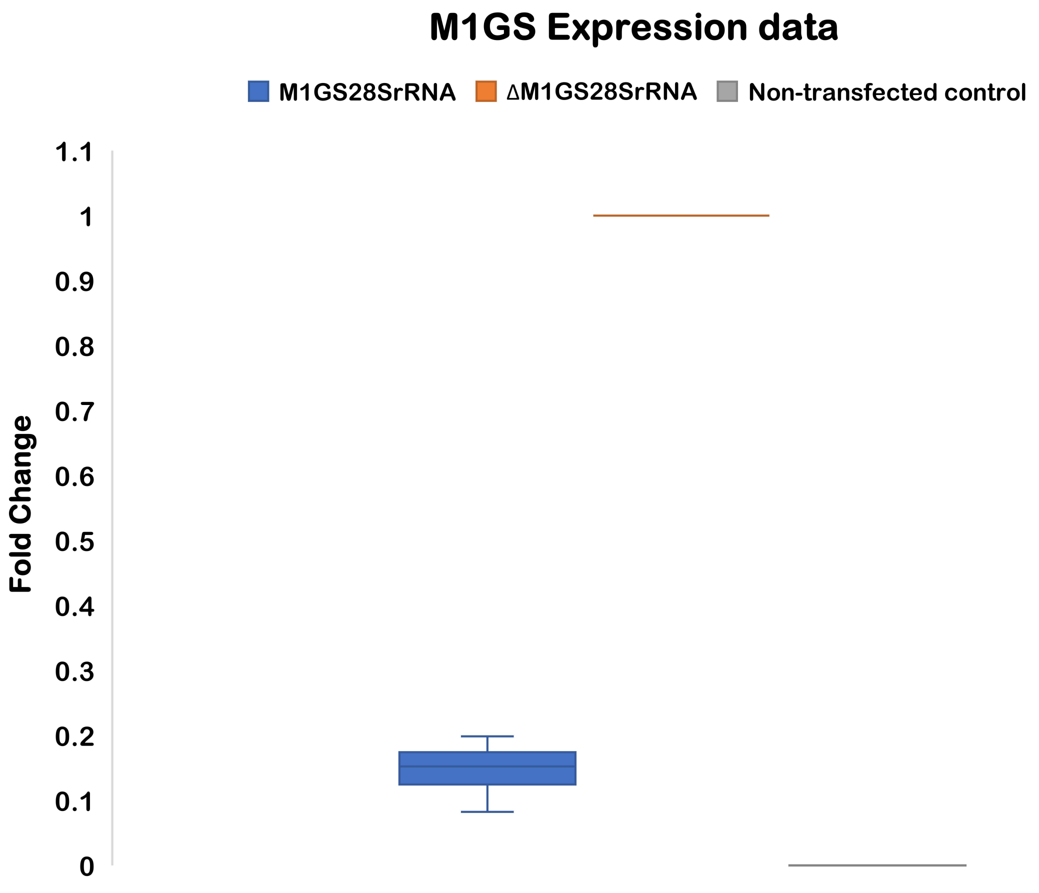


**Figure S7:** A box plot analysis of qPCR data depicts active and inactive M1GS expression levels in the human A549 cell line. M1GS expression levels are normalized to those of ΔM1GS.

**A. Sequence of thymidine kinase (TK) of human herpes simplex virus (HSV-1) – Liu & Altman, 1996**

Target RNA : 5'-GUCUGCGUUCGACCAGGCU-3'

>OR232881.1 Human alphaherpesvirus 1 isolate SP1.UL23.14 thymidine kinase gene, complete cds

```
ATGGCTTCGTACCCCTGCCATCAACACGCGTCTGCGTTCGACCAGGCTGCGCGTTCTCGCGGCCATAGCAACCGACGTACGG
CGTTGCGCCCTCGCCGGCAGCAAGAAGCCACGGAAGTCCGCCTGGAGCAGAAAATGCCCACGCTACTGCGGGTTTATATAGA
CGGTCCTCACGGGATGGGGAAAACCACCACCACGCAACTGCTGGTGGCCCTGGGTTCGCGCGACGATATCGTCTACGTACCC
GAGCCGATGACTTACTGGCAGGTGCTGGGGGCTTCCGAGACAATCGCGAACATCTACACCACACAACACCGCCTCGACCAGG
GTGAGATATCGGCCGGGGACGCGGCGGTGGTAATGACAAGCGCCCAGATAACAATGGGCATGCCTTATGCCGTGACCGACGC
CGTTCTGGCTCCTCATATCGGGGGGGAGGCTGGGAGCTCACATGCCCCGCCCCCGGCCCTCACCCTCATCTTCGACCGCCAT
CCCATCGCCGCCCTCCTGTGCTACCCGGCCGCGCGATACCTTATGGGCAGCATGACCCCCCAGGCCGTGCTGGCGTTCGTGG
CCCTCATCCCGCCGACCTTGCCCGGCACAAACATCGTGTTGGGGGCCCTTCCGGAGGACAGACACATCGACCGCCTGGCCAA
ACGCCAGCGCCCCGGCGAGCGGCTTGACCTGGCTATGCTGGCCGCGATTCGCCGCGTTTACGGGCTGCTTGCCAATACGGTG
CGGTATCTGCAGGGCGGCGGGTCGTGGCGGGAGGATTGGGGACAGCTTTCGGGGACGGCCGTGCCGCCCCAGGGTGCCGAGC
CCCAGACCAACGCGGGCCCACGACCCCATATCGGGGACACGTTATTTACCCTGTTTCGGGCCCCCGAGTTGCTGGCCCCCAA
CGGCGACCTGTATAACGTGTTTGCCTGGGCCTTGGACGTCTTGGCCAAACGCCTCCGTCCCATGCACGTCTTTATCCTGGAT
TACGACCAATCGCCCGCCGGCTGCCGGGACGCCCTGCTGCAACTTACCTCCGGGATGGTCCAGACCCACGTCACCACCCCAG
GCTCCATACCGACGATCTGCGACCTGGCGCGCACGTTTGCCCGGGAGATGGGGGAGGCTAACTGA
```

**B. Screenshot depicting the list of predicted M1GS target sites (Outputs)**

Results:

Target sites with C at -1

| Position of -1 site in the target RNA | Target site sequence (from +1 position) | % GC content |
|---|---|---|
| 8 | 5'-GUACCCCUGCCAUCAACAC-3' | 57.9 |
| 29 | 5'-GUCUGCGUUCGACCAGGCU-3' | 63.2 |
| 39 | 5'-GACCAGGCUGCGCGUUCUC-3' | 68.4 |
| 52 | 5'-GUUCUCGCGGCCAUAGCAA-3' | 57.9 |
| 76 | 5'-GUACGGCGUUGCGCCCUCG-3' | 73.7 |
| 83 | 5'-GUUGCGCCCUCGCCGGCAG-3' | 78.9 |
| 113 | 5'-GGAAGUCCGCCUGGAGCAG-3' | 68.4 |
| 321 | 5'-GACCAGGGUGAGAUAUCGG-3' | 57.9 |
| 405 | 5'-GACGCCGUUCUGGCUCCUC-3' | 68.4 |
| 566 | 5'-GUUCGUGGCCCUCAUCCCG-3' | 68.4 |
| 570 | 5'-GUGGCCCUCAUCCCGCCGA-3' | 73.7 |
| 711 | 5'-GUUUACGGGCUGCUUGCCA-3' | 57.9 |
| 766 | 5'-GGGAGGAUUGGGGACAGCU-3' | 63.2 |
| 788 | 5'-GGGGACGGCCGUGCCGCCC-3' | 89.5 |
| 803 | 5'-GCCCCAGGGUGCCGAGCCC-3' | 84.2 |
| 1014 | 5'-GCCCUGCUGCAACUUACCU-3' | 57.9 |
| 1094 | 5'-GCGCACGUUUGCCCGGGAG-3' | 73.7 |
| 1108 | 5'-GGGAGAUGGGGGAGGCUAA-3' | 63.2 |

**Figure S8: M1GS target site prediction for thymidine kinase of HSV-1. (A)** Sequence of thymidine kinase (TK). The sequence highlighted in yellow indicates the target sequence used to design GS. **(B)** A screenshot shows a list (output) of M1GS target sites predicted for the TK RNA. The target site enclosed in a red rectangle in the diagram was the GS used in that study (Liu & Altman, 1996).

### A. Sequence of 5' untranslated region (UTR) of hepatitis C virus (HCV) – Mao et al., 2014

Target RNA: 5'-GCAGAAAGCGUCU-3'

>AB828701.1 Hepatitis C virus subtype 1b genomic RNA, complete genome, strain: Pt1-98_Q2

(ONLY FIRST 341 NUCLEOTIDES)

GCCAGCCCCCGATTGGGGGCGACACTCCACCATAGATCACTCCCCTGTGAGGAACTACTGTCTTCACGCAGAAAGCGTCTAG
CCATGGCGTTAGTATGAGTGTCGTGCAGCCTCCAGGACCCCCCCTCCCGGGAGAGCCATAGTGGTCTGCGGAACCGGTGAGT
ACACCGGAATTGCCAGGACGACCGGGTCCTTTCTTGGATCAACCCGCTCAATGCCTGGAGATTTGGGCGTGCCCCCGCGAGA
CTGCTAGCCGAGTAGTGTTGGGTCGCGAAAGGCCTTGTGGTACTGCCTGATAGGGTGCTTGCGAGTGCCCCGGGAGGTCTCG
TAGACCGTGCACC

### B. Secondary structure predicted by mfold

Input

DNA or RNA sequence of the target RNA

GCCAGCCCCCGAUUGGGGGCGACACUCCACCAUAGAUCACUCCCCUGUGAGGAACUACUGUCUU
CACGCAGAAAGCGUCUAGCCAUGGCGUUAGUAUGAGUGUCGUGCAGCCUCCAGGACCCCCCCUC
CCGGGAGAGCCAUAGUGGUCUGCGGAACCGGUGAGUACACCGGAAUUGCCAGGACGACCGGGU

NOTE:

If you want to use your own secondary structure, please paste the secondary structure of your target sequence in the vienna format. If you do not have one, a secondary structure predicted by the RNAFold Software will be used.

Example: ((....))((((((((.........)))).....)))......((....())....)

Secondary structure of the target RNA in Vienna format (Optional)

....((((((....))))))((((((((..................(((((((.(....))))))))......((((((........))))))......)))))))(((((..((((((((......(((....))).
((((.....((.((((((((..((((((.....))))))..)))))..)))).))..((((((.......))))))......))))....))))))))..(..((((((((.....(((.(((((((......)))))).)))
((((.......))))....)))))).)))..)..(((..((....))..)))......((((......)))).)))))).

Nucleotide at -1 position: C (on), U, A, G

Nucleotide at +1 position: G (on), A

CCA at 3' position of M1GS unpaired

Unpaired nucleotide at -1 position according to the predicted structure of the target RNA

Desirable length of the target RNA from -1 position that is predicted to exist in a single-stranded form: 6

Length of the Guide Sequence (GS): 13

Results:

Target sites with C at -1

| Position of -1 site in the target RNA | Target site sequence (from +1 position) | % GC content |
|---|---|---|
| 67 | 5'-GCAGAAAGCGUCU-3' | 53.8 |
| 270 | 5'-GCGAAAGGCCUUG-3' | 61.5 |

### C. Secondary structure predicted by RNAfold

Input

DNA or RNA sequence of the target RNA

GCCAGCCCCCGAUUGGGGGCGACACUCCACCAUAGAUCACUCCCCUGUGAGGAACUACUGUCUU
CACGCAGAAAGCGUCUAGCCAUGGCGUUAGUAUGAGUGUCGUGCAGCCUCCAGGACCCCCCCUC
CCGGGAGAGCCAUAGUGGUCUGCGGAACCGGUGAGUACACCGGAAUUGCCAGGACGACCGGGU

NOTE:

If you want to use your own secondary structure, please paste the secondary structure of your target sequence in the vienna format. If you do not have one, a secondary structure predicted by the RNAFold Software will be used.

Example: ((....))((((((((.........)))).....)))......((....())....)

Secondary structure of the target RNA in Vienna format (Optional)

....((((((....))))))((((((((..................(((((((.((....)))))))))......((((((........))))))......)))))))(((((...((..(((((.(((.....((((((.
((((((.(((((.((..(((((...((((.((.(((((.(((((..((.(((((((((.......)))....)))).))....))))).))))).)).)))))..)).(((.
((((((......)))))).))))))..))..)))))))))))).)).))))...(((((((.....))))))).))).)))))....))...))))).

Nucleotide at -1 position: C (on), U, A, G

Nucleotide at +1 position: G (on), A

CCA at 3' position of M1GS unpaired

Unpaired nucleotide at -1 position according to the predicted structure of the target RNA

Desirable length of the target RNA from -1 position that is predicted to exist in a single-stranded form: 6

Length of the Guide Sequence (GS): 13

Results:

No target sites found with C at -1 with the set parameters

### D. Secondary structure predicted by RNAfold

Input

DNA or RNA sequence of the target RNA

CCUUUCUUGGAUCAACCCGCUCAAUGCCUGGAGAUUUGGGCGUGCCCCCGCGAGACUGCUAGC
CGAGUAGUGUUGGGUCGCGAAAGGCCUUGUGGUACUGCCUGAUAGGGUGCUUGCGAGUGCCCC
GGGAGGUCUCGUAGACCGUGCACC

NOTE:

If you want to use your own secondary structure, please paste the secondary structure of your target sequence in the vienna format. If you do not have one, a secondary structure predicted by the RNAFold Software will be used.

Example: ((....))((((((((.........)))).....)))......((....())....)

Secondary structure of the target RNA in Vienna format (Optional)

Enter sequence here

Nucleotide at -1 position: C (on), U, A, G

Nucleotide at +1 position: G (on), A

CCA at 3' position of M1GS unpaired

Unpaired nucleotide at -1 position according to the predicted structure of the target RNA

Desirable length of the target RNA from -1 position that is predicted to exist in a single-stranded form: 0

Length of the Guide Sequence (GS): 13

Results:

Target sites with C at -1

| Position of -1 site in the target RNA | Target site sequence (from +1 position) | % GC content |
|---|---|---|
| 10 | 5'-GAUUGGGGGCGAC-3' | 69.2 |
| 20 | 5'-GACACUCCACCAU-3' | 53.8 |
| 67 | 5'-GCAGAAAGCGUCU-3' | 53.8 |
| 76 | 5'-GUCUAGCCAUGGC-3' | 61.5 |
| 89 | 5'-GUUAGUAUGAGUG-3' | 38.5 |
| 104 | 5'-GUGCAGCCUCCAG-3' | 69.2 |
| 130 | 5'-GGGAGAGCCAUAG-3' | 61.5 |
| 151 | 5'-GGAACCGGUGAGU-3' | 61.5 |
| 157 | 5'-GGUGAGUACACCG-3' | 61.5 |
| 169 | 5'-GGAAUUGCCAGGA-3' | 53.8 |
| 183 | 5'-GACCGGGUCCUUU-3' | 61.5 |
| 187 | 5'-GGGUCCUUUCUUG-3' | 53.8 |
| 209 | 5'-GCUCAAUGCCUGG-3' | 61.5 |
| 232 | 5'-GUGCCCCCGCGAG-3' | 84.6 |
| 240 | 5'-GCGAGACUGCUAG-3' | 61.5 |
| 242 | 5'-GAGACUGCUAGCC-3' | 61.5 |
| 255 | 5'-GAGUAGUGUUGGG-3' | 53.8 |
| 270 | 5'-GCGAAAGGCCUUG-3' | 61.5 |
| 272 | 5'-GAAAGGCCUUGUG-3' | 53.8 |
| 308 | 5'-GAGUGCCCCGGGA-3' | 76.9 |
| 317 | 5'-GGGAGGUCUCGUA-3' | 61.5 |
| 327 | 5'-GUAGACCGUGCAC-3' | 61.5 |
| 334 | 5'-GUGCACC-3' | 71.4 |

**Figure S9: M1GS target site prediction for hepatitis C subtype 1b genomic RNA.**
(A) The hepatitis C subtype 1b genomic RNA sequence used in the study and the sequence highlighted in yellow indicate the target sequence used to design GS. Screenshots show the output when the secondary structure for the target RNA was predicted by (B) Mfold software and the resultant prediction data in the Vienna format was provided as an input for the tool, (B&C) RNAfold software for different inputs. In (B-D), The target site enclosed in a red rectangle was used for designing GS for cleaving the target RNA in the study (Mao et al. 2014)

| *Primer* | *Forward (5’-3’)* | *Reverse (5’-3’)* |
|---|---|---|
| 28S rRNA | ATGTAGGTAAGGGAAGTCGGCAAGCC | ACCGACCCAGCCCTTAGAGCCAATC |
| M1GS_HH | TAACAGTCCGTGGCACGGTAAACTCC | CAGGACATCGGGTTTCAGATGTCGAC |
| Active M1GS | TAACAGTCCGTGGCACGGTAAACTCC | GACCGATAAGCCGGGTTCTGT |
| Inactive M1GS | TAACAGTCCGTGGCACGGTAAACTCC | GTGAAACTGACCGAGATACATCTCGCTAG |
| GAPDH | CTCCTGCACCACCAACTGCT | GGGCCATCCACAGTCTTCTG |

**Figure S10: List of primers used in the study**
